# Episodic events are flexibility encoded in both integrated and separated neural representations

**DOI:** 10.1101/2025.04.30.651466

**Authors:** Zhenghao Liu, Mikael Johansson, Inês Bramão

## Abstract

This study investigates how the brain encodes episodic events to support diverse memory functions. Thirty-six participants viewed movies simulating real-life interactions while EEG was recorded. They first watched movies featuring two characters (AB), followed by scenes where one original character interacted with a new one (BC). Memory was assessed for direct (AB and BC), indirect (AC) associations, and source memory about whether two characters had appeared together. Source memory was more accurate when participants correctly inferred indirect associations. Time-resolved representational similarity analysis (RSA) of EEG data revealed both neural pattern similarities and dissimilarities during AB and BC encoding. Pattern similarities, indicating memory integration across episodes, predicted successful indirect associations. Conversely, dissimilarities predicted accurate source memory, indicating the preservation of distinct memory traces. These findings suggest that episodic events are flexibly encoded through both integrated and segregated neural representations, enabling the brain to support multiple memory functions depending on task demands.

## INTRODUCTION

Episodic memory enables the adaptive retrieval of commonalities and unique details across overlapping life events. For example, encountering a woman in the park with your colleague’s daughter may lead to the formation of an integrated memory representation connecting the woman and your colleague. At the same time, you may retain a distinct representation of this specific encounter to later discuss with your colleague the circumstances of meeting her daughter. This study investigates how the brain simultaneously supports both the integration across related events and the preservation of event-specific details.

Functional magnetic resonance imaging (fMRI) research has examined how the brain represents memories for related events by analysing the neural pattern similarities associated with different events. Some memory models suggested that the hippocampus encodes related events into distinct memory representations to minimise interference^1–3^. However, more recent evidence suggests that the hippocampus also supports the integration of related events^4,5^, facilitating the inference of indirect associations between elements encountered in separated episodes^6,7^. To reconcile these apparently conflicting findings, it has been proposed that hippocampal function varies along its axis, with the anterior hippocampus supporting the formation of integrated memory representations and the posterior hippocampus forming distinct, event-specific memory representations^8,9^.

Despite evidence for the formation of both integrated and separated memory representations, it remains unclear whether these representations can coexist for the same event or if integration compromises event-specific detail. Some findings suggest that memory integration comes at the cost of diminished episodic detail. In particular, Carpenter et al.^10,11^ demonstrated that integrated representations are associated with reduced memory for episodic details, indicating a trade-off between memory integration and event-specific detail. This raises the possibility that the brain cannot simultaneously keep both types of representations. Nevertheless, this trade-off has not been consistently replicated, and some studies even report a positive association between memory integration and episodic detail memory^12–14^, suggesting that integrated and separated representations may coexist^15^.

It is plausible that the brain maintains both types of representations to flexibly support different memory functions^8,15^. Integrated representations may encode generalised information, facilitate inferences across events, and enable the seamless retrieval of overlapping memories. In contrast, separated representations preserve event-specific features, reduce interference, and support the retrieval of unique event details. Previous fMRI studies have provided evidence that integrated and separated neural representations emerge in different hippocampal subregions^9^. However, it remains unclear how these representations are formed during real-time encoding when new episodic events overlap with past experiences. Specifically, little is known about how the brain simultaneously integrates novel information into existing memory structures while preserving the distinctiveness of individual events. Do integration and separation processes dynamically interact during encoding? How does the brain determine which aspects of an experience should be merged and which should remain distinct? To address this gap, we employed time-resolved representational similarity analysis (RSA) of electroencephalographic (EEG) recordings to track the dynamic formation of neural patterns during real-time encoding of related events. This approach allowed us to examine the emergence of integrated and separated representations as they develop in real-time and to investigate how they support different memory functions. Furthermore, by exploring how shared and novel information across related events contributes to the formation of these neural representations, we provide insight into the mechanisms that govern the interplay between memory integration and separation during episodic encoding.

We used a naturalistic paradigm, adapted from the classic associative inference memory paradigm^5,16,17^, in which participants encode videos that simulate real-life events, allowing us to capture the complexity of memory formation in an ecologically valid setting. Participants encoded movies featuring real-world interactions between *Sim* characters (https://www.ea.com/games/the-sims/the-sims-4). First, they encoded AB movies, in which a Sim A interacted with a Sim B. Later, they encoded BC movies, where a new Sim C interacted with the previously seen Sim B. A control condition featured interactions between two novel Sims (XY movies). During retrieval, participants completed memory tests for both direct (i.e., AB, BC, XY) and indirect (i.e., AC) associations. These tests were interleaved, with the constraint that AC associations were tested before their corresponding AB and BC associations. Each association memory test was followed by a source memory task, where participants indicated if the two characters had directly interacted. Participants also rated their confidence (see **Figure 1A**). High AC performance was considered an indicator of the formation of integrated representations across events^5,16,17^, while high source memory performance reflected preserved event-specific representations (see also^10,18^). An additional surprise memory test for episodic details (context and character clothing) showed no systematic effects of AC retrieval and is reported in the supplementary material (see **Supplementary Note 1**).

**Figure 1.**
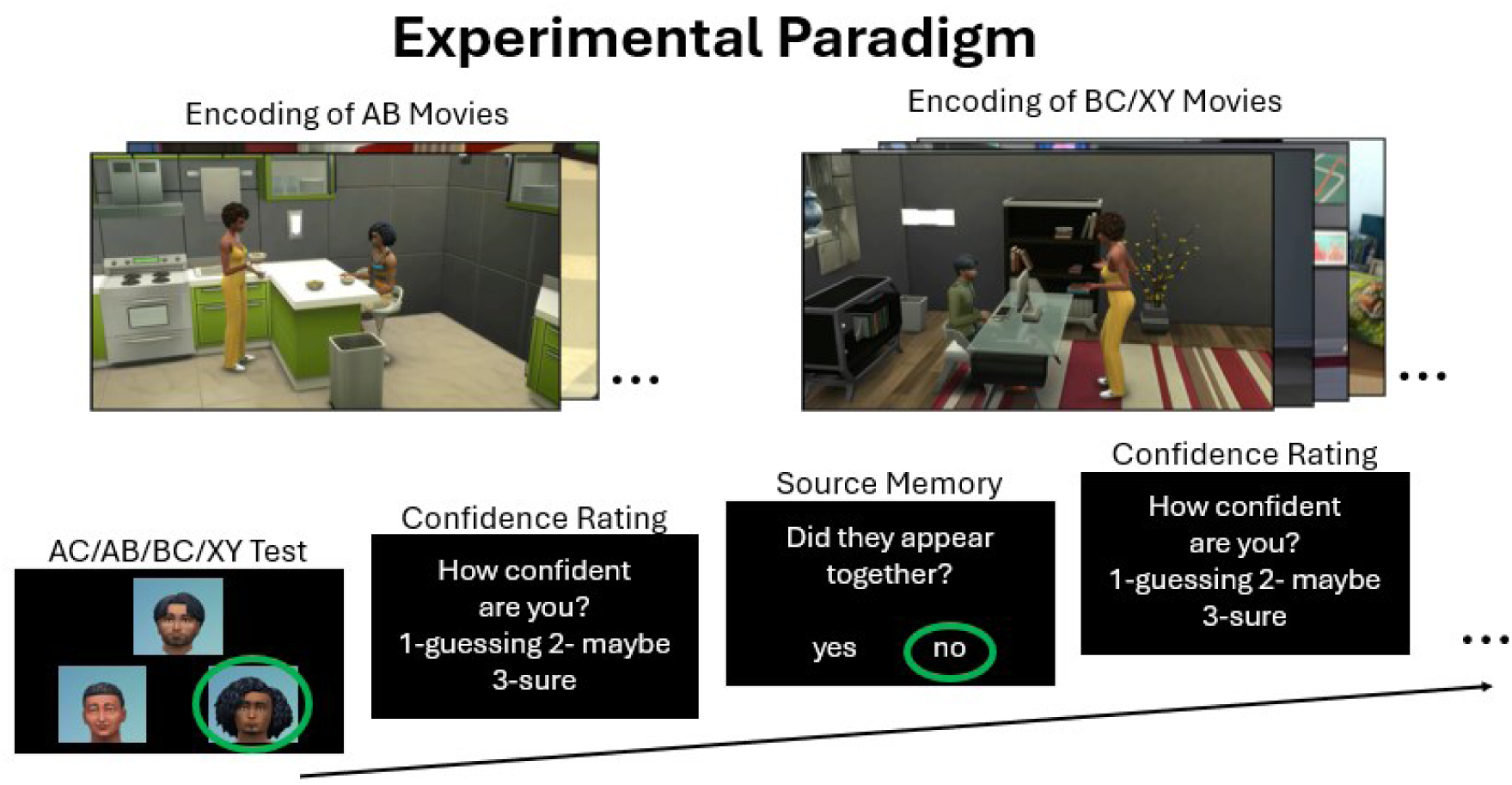
Overview of the experimental paradigm. Participants first encoded AB movies in which they encountered two characters. Later, they encoded BC movies, where a new character interacted with a previously seen one. XY movies were also presented with two completely new characters and serve as a control condition. Each movie was presented five times during encoding. At retrieval, participants completed memory tests for direct (AB, BC and XY) and indirect associations (AC). The retrieval test order was randomized, with the constraint that AC associations were always tested before their corresponding AB and BC associations. Following each memory test, participants completed a source memory test to indicate if the two characters had been seen together. Confidence ratings were also registered.

EEG data were recorded throughout the experiment. We applied RSA to the EEG data during the encoding of AB, BC and XY movies, which allowed us to track the formation of the integrated and separated memory representations (see **Figure 2**). Integration was expected to be reflected in greater neural similarity between AB and BC movies, whereas separation indicated by neural dissimilarities between AB and BC movies relative to control conditions. We further hypothesised that neural similarities predict AC retrieval, while neural dissimilarities predict source memory. Specifically, if integrated and separated representations coexist, successful AC retrieval should be accompanied by better source memory, indicating that both types of representations, supporting different memory functions, were encoded for the same events. Conversely, if integration disrupts event-specific details, successful AC retrieval should impair source memory, and the neural patterns predictive of AC retrieval would associate with poorer source memory performance.

**Figure 2.**
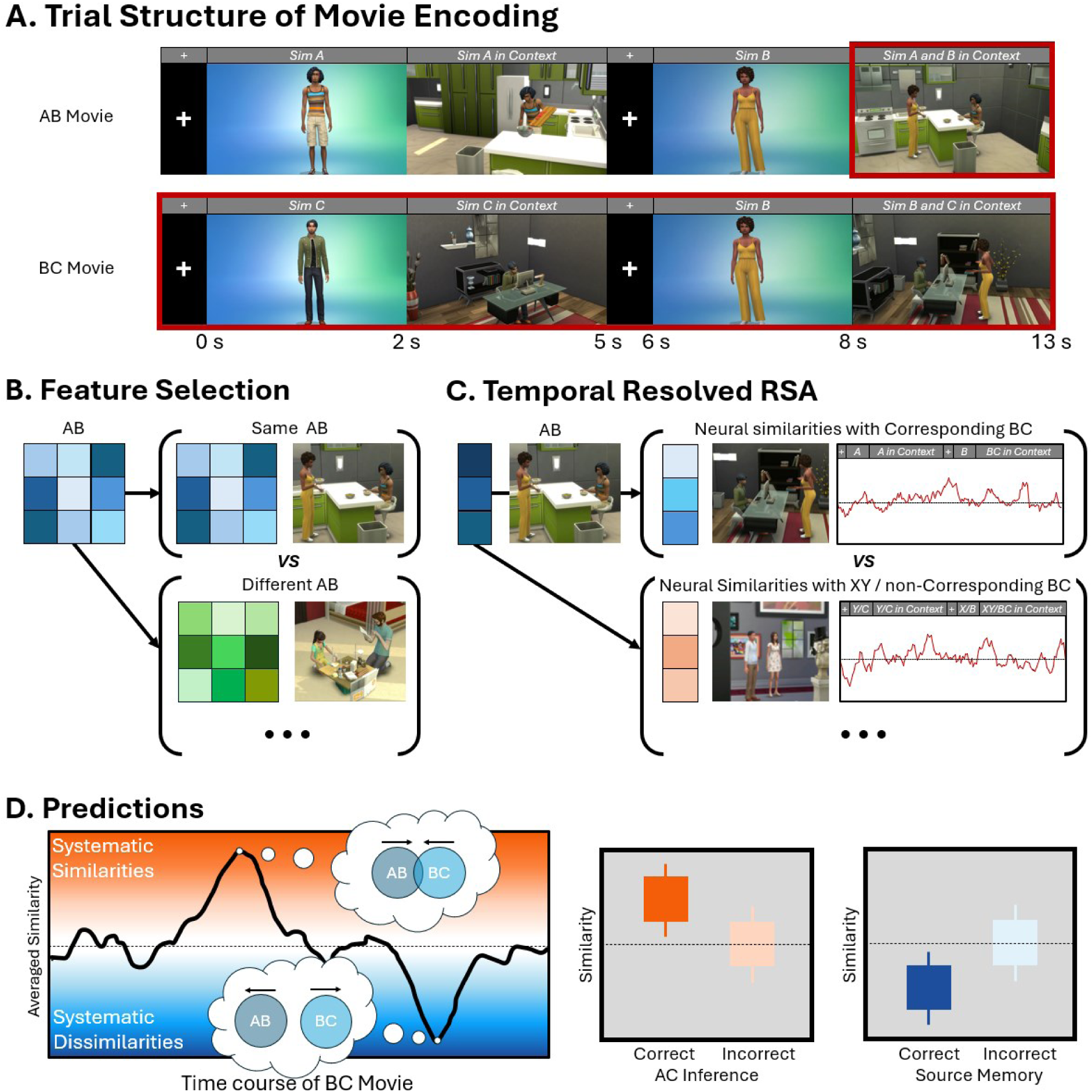
Analytical pipeline and predictions of the present study. (A) Trial structure. Each movie began with the display of Sim A/C/Y for two seconds, followed by a three-second animation of this Sim in context. Afterward, a one-second fixation cross was presented, followed by the presentation of the Sim B/X for two seconds. The movie ended with a five-second animation of the two Sims AB/BC/XY interacting in a context. We measured the neural representational similarity of the *Sim A and B in Context* of AB movie (outlined in red) and each time point of the corresponding BC movie (outlined in red). (B) Feature Selection. To identify the time-frequency features that were sensitive to the content of the AB movies, we compared the wavelet coherence between different repetitions of the same AB movie against the coherence between different AB movies. (C) A time-resolved neural representational similarity analysis was performed by correlating the representative features of the AB movie with the patterns of the corresponding BC movie across the entire encoding period. Systematic similarities and dissimilarities were estimated by contrasting the neural patterns similarities between AB and corresponding BC movie against two baselines: 1) neural similarities between AB and XY movies, and 2) neural similarities between AB and non-corresponding BC movies. (D) Systematic similarities, indicative of memory integration should predict of AC retrieval performance, whereas systematic dissimilarities, indicative of memory separation, should predict source memory performance.

## RESULTS

### Behavioural Results

#### Associative Memory Tests

Memory performance for direct and indirect associations are summarized in **Figure 3A**. We contrasted memory performance across the four association types (AC vs. AB vs. BC vs. XY) with linear mixed-effects models. This analysis showed significant main effects of association type on accuracy (*F*(3, 3420) = 48.068, *p* < .001, 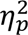 = .04), response time (*F*(3, 2587) = 71.314, *p* < .001, 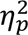 = .08) and confidence (*F*(3, 2584) = 11.921, *p* < .001, 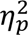 = .01). Post-hoc Tukey-corrected comparisons showed that retrieval of AC associations was associated with significantly lower accuracy, slower response times, and reduced confidence compared with all the direct associations (*p*s < .001), except that the lower confidence of AC compared to XY did not reach statistical significance (*t*(2590) = −2.427, *p* = .072, *D* = −0.045).

**Figure 3.**
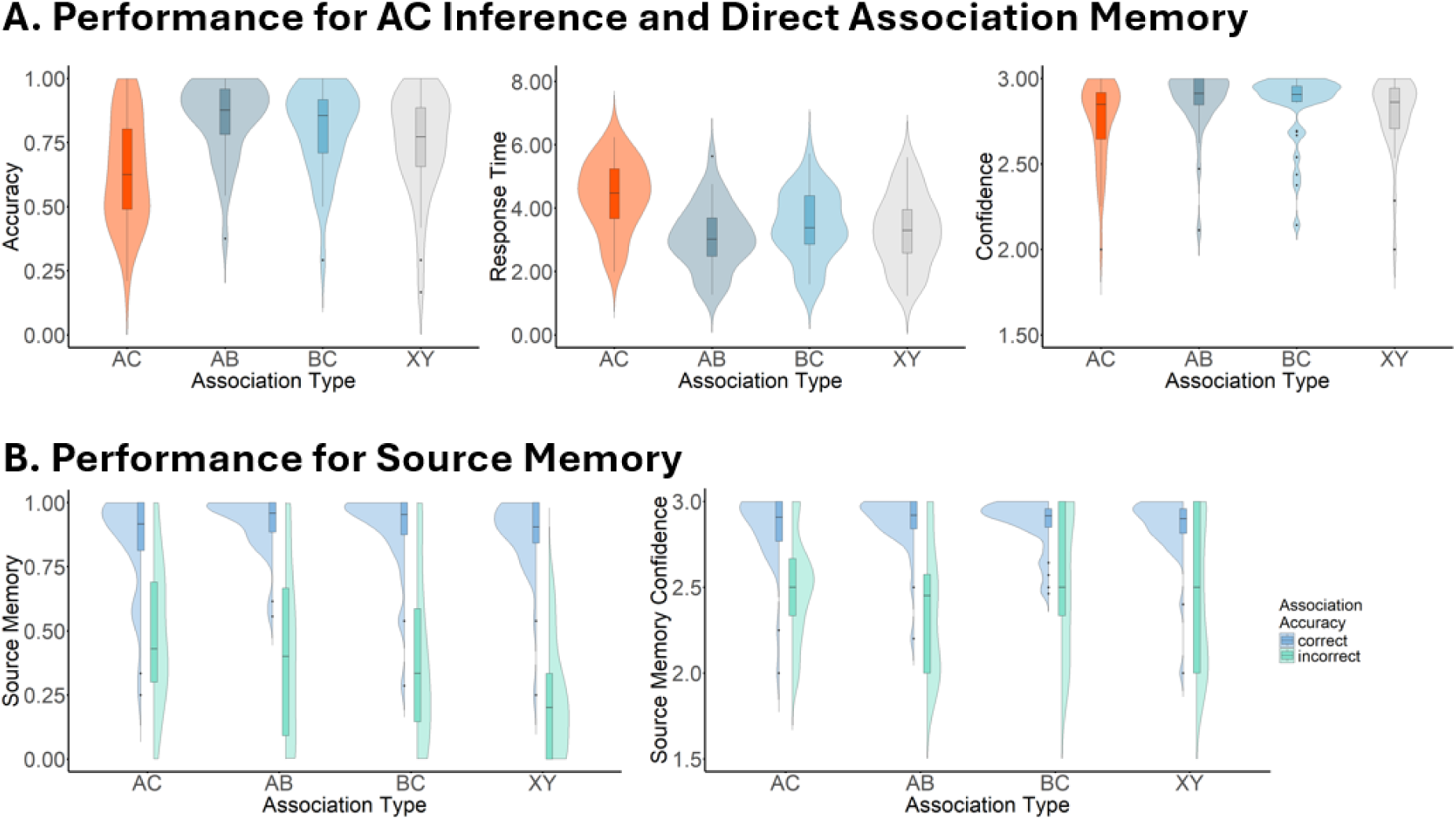
Behavioural result summary. (A) Accuracy, response time and confidence for the memory tests (AC vs AB vs BC vs XY). Violin plots show the sample distribution, and the box plots indicate the means and quartiles for each measure. (B) Accuracy and confidence for the source memory tests of each association type (AC vs AB vs BC vs XY) and retrieval accuracy (Correct vs. Incorrect) of the corresponding association test. Raincloud plots show the distribution sample of the data, with the box plots indicating the means and quartiles.

Furthermore, XY associations were retrieved less accurately than both AB (*t*(3423) = 5.37, *p* < .001, *D* = 0.101) and BC (*t*(3423) = 3.15, *p* = .009, *D* = 0.059) associations, while no difference was found between the AB and BC retrieval accuracy (*t*(3423) = 2.22, *p* = .118, *D* = 0.042). In terms of response times, BC was retrieved slower than both AB (*t*(2590) = −4.85, *p* < .001, *D* = −0.150) and XY (*t*(2590) = −2.98, *p* = .015, *D* = −0.096), but no differences were observed between AB and XY associations (*t*(2591) = −1.726, *p* = .310, *D* = −0.054). Finally, AB associations were retrieved more confidently than XY associations (*t*(2591) = 3.11, *p* = .010, *D* = 0.054), while the confidence of BC was not significantly different from either AB or XY retrieval (*t*s < 2.29, *p*s > .101).

### Source Memory Test

To evaluate how source memory performance varied as a function of retrieval accuracy across the different associations, we used linear mixed-effects models with Association Type (AC vs. AB vs. BC vs. XY) and Retrieval Accuracy (Correct vs. Incorrect) as factors, see **Figure 3B**. The results show a significant main effect of Association Type for accuracy (*F*(3, 36) = 5.329, *p* = .004, 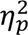 = .31) but not for confidence (*F*(3, 126) = 1.947, *p* = .125, 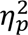 = .04), showing worse source memory for XY compared with both AC (*t*(33) = −3.362, *p* = .010, *D* = −0.083) and AB associations (*t*(30) = −3.236, *p* = .015, *D* = −0.108). All the other contrasts were non-significant (*p*s > .091). The main effect of Retrieval Accuracy was also significant for both accuracy (*F*(1, 26) = 293.667, *p* < .001, 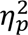 = .92) and confidence (*F*(1, 2656) = 282.903, *p* < .001, 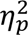 = .10), showing that when the associations were correctly retrieved, source memory performance was higher and also associated with higher confidence ratings.

Additionally, a significant interaction between Association Type and Retrieval Accuracy was observed for both source memory accuracy (*F*(3, 34) = 11.007, *p* < .001, 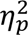 = .49) and confidence (*F*(3, 2523) = 2.802, *p* = .039, 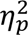 = 3.32e-3). Post-hoc tests confirmed that higher source memory accuracy and confidence were consistently associated with correct association retrieval (*p*s < .001), suggesting that the observed interactions were primarily driven by the effect size differences across Association Types. Monte Carlo sampling further revealed that the differences in source memory accuracy between correct and incorrect retrieved associations was largest for XY associations (*p*s < .001), whereas the other associations showed comparable levels of this difference (*p*s > .128). On the other hand, the effect size of retrieval accuracy on source memory confidence was smallest for BC associations (*p*s < .050), while the effect size was comparable across the other association types (*p*s > .346). In summary, the behavioural results show that source memory performance is positively associated with AC memory performance, supporting the idea that related events may be simultaneously encoded in both integrated and separated memory representations. These different memory representations may facilitate distinct memory functions and support different mnemonic task demands.

## EEG Results

### Feature Selection Reveals Time-Frequency Signatures of AB Movie Content

The EEG data collected during the encoding of each AB movie, was transformed into a time-frequency representation (TFR), for use in the neural representational similarity analysis (RSA). Before conducting the RSA, we identified representative time-frequency features that reliably captured the contents of AB movie. Those features should remain stable across repetitions of the same AB movie and are distinct across different AB movies.

To achieve this, we applied a wavelet coherence-based approach^19^. Specifically, we computed the coherence across the five repetitions of each AB movie (within-AB coherence) and contrasted it with the coherence between the different AB movies (between-AB coherence; see **Figure 2B**). Time-frequency features exhibiting significantly higher within-AB coherence than between-AB coherence were considered sensitive to the context of AB and selected for the RSA. **Figure 4** presents the results of this feature selection, averaged across all AB movies. A cluster-based permutation test revealed four significant clusters. The first cluster emerged immediately after the movie onset, between 0-0.9 seconds, corresponding to the initial presentation of Sim A (*t*-cluster = 1.229e+4, *p* = .003). A second cluster appeared between 2-3 seconds, capturing the representation of Sim A within its context (*t*-cluster = 3.344e+4, *p* < .001). The third cluster, occurring between 6-7.1 seconds, reflected the distinct neural representation of Sim B (*t*-cluster = 1.821e+4, *p* = .001). Finally, the fourth cluster, observed between 8-9.2 seconds, corresponded to when both Sim A and B appeared within a context, capturing the full event representation (*t*-cluster = 3.251e+4, *p* < .001).

**Figure 4.**
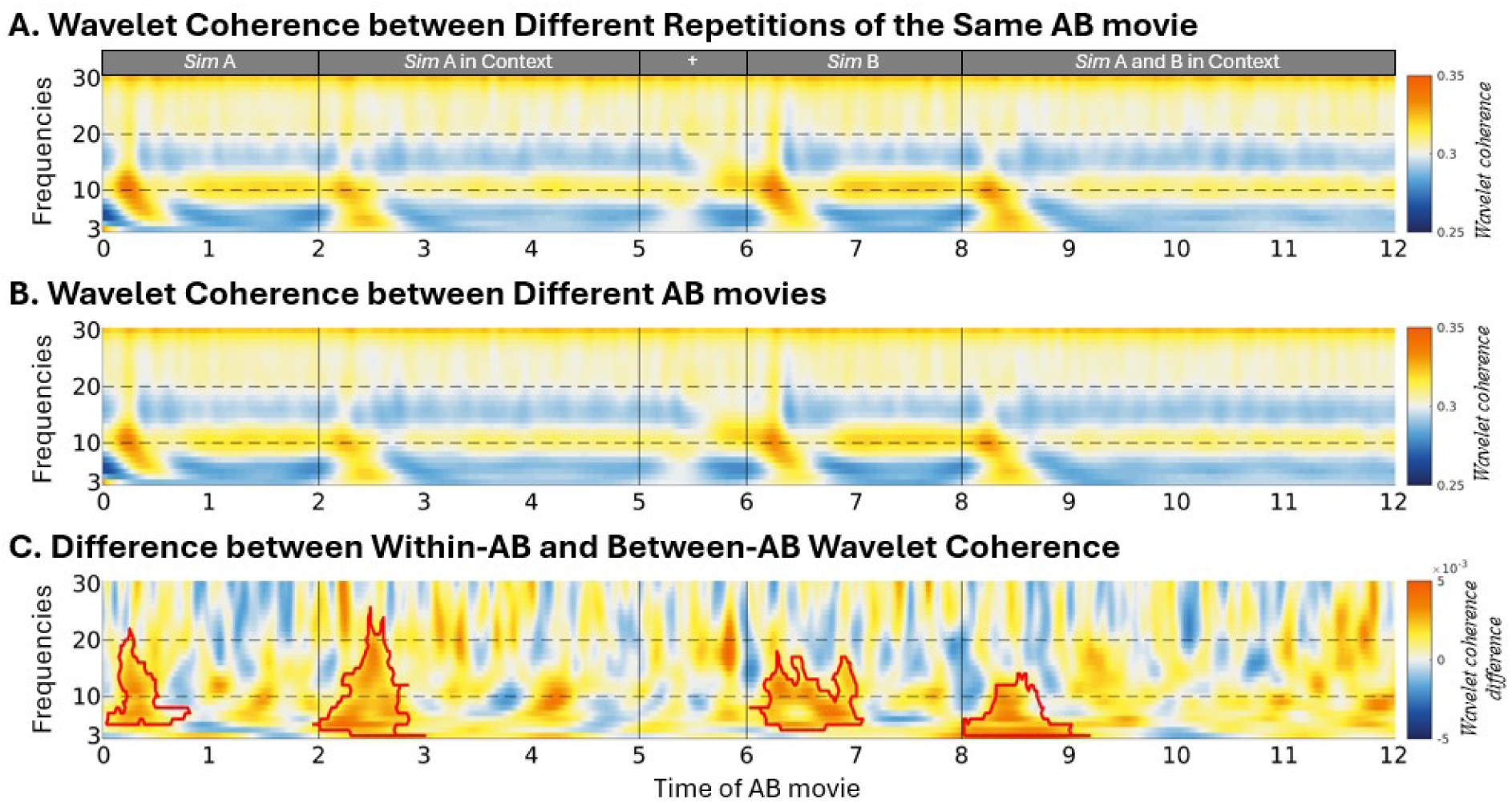
Results for the within-AB wavelet coherence (A), between-AB wavelet coherence (B) and their differences (C). Significant clusters, marked by red edges, indicate the time-frequency features that were stable across the five AB repetitions and sensitive to different AB movie content. The identified features were used as a template to extract the power values of the AB movies, which were used in the subsequent representational similarity analysis.

These AB movie-specific features were then used as a template to extract the power values from the TFR of AB trials, which were then used to estimate their representational similarity with the TFR of the XY and BC trials. Our analysis focused on the similarity between the time window corresponding to the Sim A and B interaction (i.e., the segment of *Sim A and B Context*) and each time point of the XY and BC movies encoding. This segment was chosen as it encapsulates the complete event structure, including both Sims, their actions, and the context (See **Figure 2A**). Additional analyses investigating the representational similarity between the other AB movie segments and XY and BC movies are reported in **Supplementary Note 3**.

### The Neural Representations of AB and BC Movies Share both Similarities and Dissimilarities

To investigate how AB and BC movies are represented at the neural level, we conducted a representational similarity analysis (RSA). First, we extracted time-frequency features sensitive to the content of each AB movie, averaging across the five repetitions to ensure robustness. To account for potential time shifts across repetitions, we used the temporal median of these features^20^. Next, for each participant and AB movie, we correlated these extracted features with the time-frequency representation (TFR) of the corresponding BC movie at each time point during BC encoding, resulting in a time-resolved representational similarity matrix. This matrix captures the systematic similarities and dissimilarities between the neural representations of AB and the corresponding BC movie. The BC movies began with the segments involving only the novel Sim C, followed by the common Sim B. Thus, when participants were first exposed to the novel Sim C, they could not yet know which corresponding AB movie it was associated with. This may have introduced distinct patterns of similarities and dissimilarities in this initial viewing, which was therefore excluded from the analysis. The RSA analysis across the five repetitions of BC movie is reported in **Supplementary Note 2**.

To determine whether the neural representations of AB and BC movies were significantly related, we first contrasted the representational similarity between each AB movie and its corresponding BC movie against the similarity between AB movies and all XY movies. Bayesian statistical analysis revealed systematic similarities and dissimilarities in several time windows (see **Figure 5A**). Specifically, increased similarity was observed centred at 1 second (average similarity = 1.1e-2, BF_10_s > 3.166), during the segment of *Sim C*, followed by two additional peaks at 2.8 seconds (average similarity = 9.0e-3, BF_10_s > 3.151) and 3.7 seconds (average similarity = 1.2e-2, BF_10_s > 3.061), during the presentation of *Sim C in Context*. In contrast, increased dissimilarities were observed later during BC encoding, at 5.1 seconds (average similarity = −8.9e-3, BF_10_s > 3.164) and 7.4 seconds (average similarity = −1.3e-2, BF_10_s > 3.185), immediately before and during the presentation of the segment of *Sim B*.

**Figure 5.**
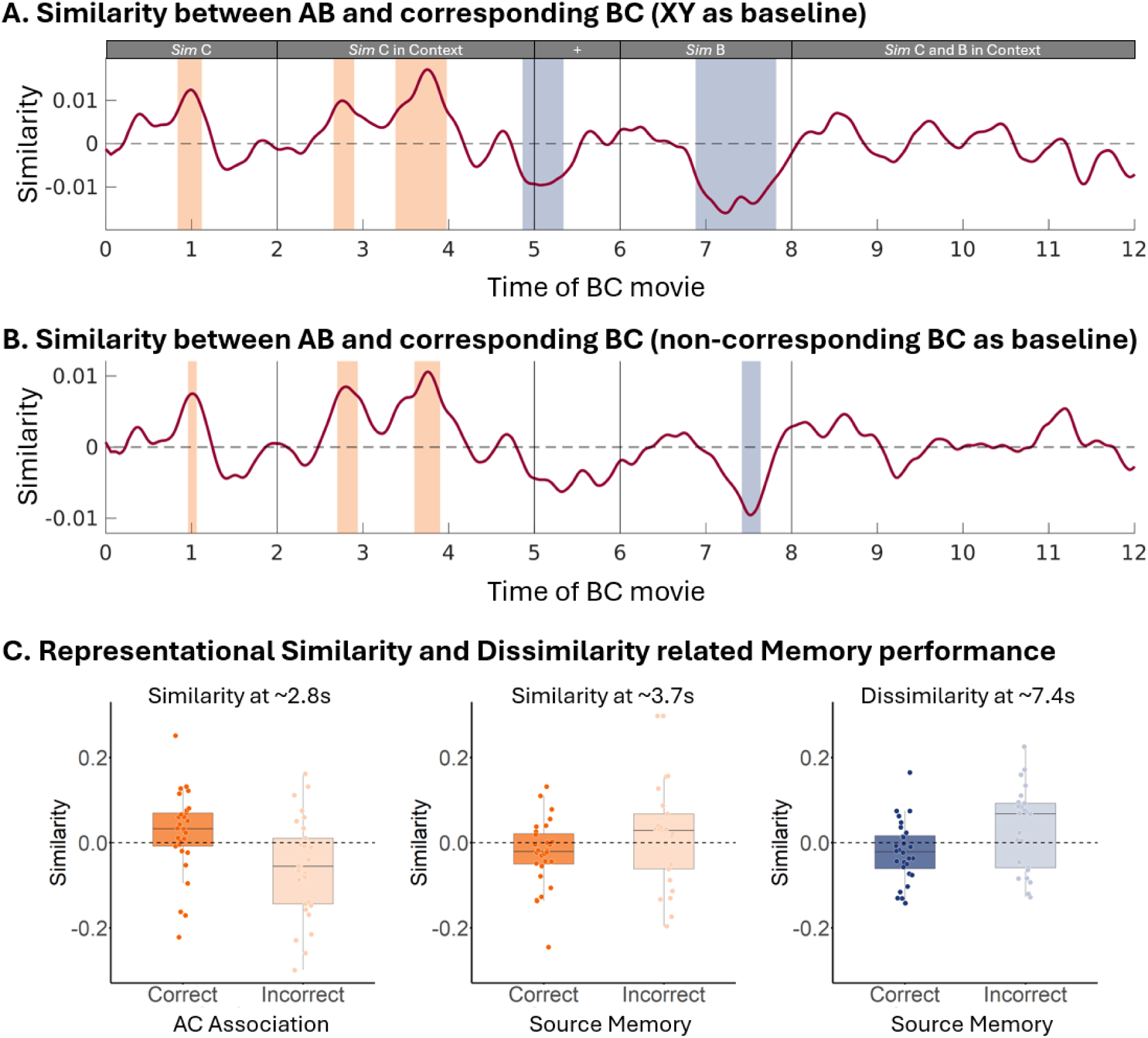
Time-resolved representational similarities and dissimilarities between AB movie and corresponding BC movie, and their relationship with memory performances. Significant time windows, identified with a Bayesian approach, are highlighted to indicate moments of increased similarity and dissimilarity. (A) Neural pattern similarities between AB and corresponding BC movie contrasted against the similarities between AB and XY movies. (B) Neural pattern similarities between AB and corresponding BC movie contrasted against the similarities between AB and non-corresponding BC movies. These findings reveal distinct phases of integration and separation in neural representations, with similarity effects emerging during Sim C encoding and dissimilarity effects occurring during the presentation of the common Sim B. (C) Representational similarities and dissimilarities for trials with correct / incorrect memories. The plotted similarities account for the effects of controlled variables and display the binary relationship between neural similarities and memory performance. Each data point reflects the average similarity per participant for correct and incorrect memory performance. The box plots indicate the mean and quartile ranges for each condition, providing a visualization of the distribution of (dis-)similarities across participants.

Next, we contrasted the similarities between the AB movie and its corresponding BC movie against the similarity evoked by the AB movie and all non-corresponding BC movies. While using the XY movies is a common baseline in the literature (for a review, see^8^), it does not entirely rule out general, content-unspecific reinstatement. The Sim B might trigger general retrieval processes in which previously encoded events are broadly reactivated in search of the corresponding AB movie. This could influence the observed representational similarity, regardless of whether the movies share a common element. Thus, to differentiate content-specific reinstatement (i.e., reinstatement of the corresponding AB movie) from the general reinstatement (i.e., broader reinstatement of multiple AB movies) we compared the similarity of AB and corresponding BC movie against the similarity between AB and all non-corresponding BC movies. This analysis allowed us to isolate neural patterns reflecting the retrieval of a specific BC movie rather than general memory reinstatement. The results of this analysis replicated the previous findings (see **Figure 5B**), with significant similarities emerging at 1 second (average similarity = 7.5e-3, BF_10_s > 3.019), 2.8 seconds (average similarity = 8.0e-3, BF_10_s > 3.188) and 3.7 seconds (average similarity = 9.2e-3, BF_10_s > 3.065) after BC movie onset. Additionally, the same dissimilarity at 7.4 seconds was observed (average similarity = - 8.8e-3, BF_10_s > 3.164), confirming that our findings are not merely the result of broader retrieval processes but instead reflected content-specific neural similarities and dissimilarities.

In **Supplementary Note 2** we report how the similarities and dissimilarities evolve across the five repetitions of BC movie encoding. Over the successive repetitions we observed incremental shifts in both similarities and dissimilarities, reflecting an adaptive process in which integrated and separated representations evolved dynamically over time.

In summary, the neural representations of AB and BC movies share both similarities and dissimilarities, revealing integrated and separated patterns. The similarities occur when the novel element, Sim C, is presented, suggesting that participants integrate overlapping elements from different experiences into unified representations. In contrast, dissimilarities arise when the common cue, Sim B, is presented, indicating a neural mechanism that preserves distinctions between similar yet distinct events, likely to minimize interference. These findings highlight the brain’s ability to flexibility balance integration and separation, which is fundamental for creating associative memory links between different experiences and for maintaining episodic memory specificity.

### Neural Similarities Relate to Associative Inference and Neural Dissimilarity Relates to Source Memory

Next, we investigated how the identified neural similarities and dissimilarities between AB and the corresponding BC movie relate to later memory performance. For each trial, we extracted the similarity between AB and corresponding BC, with the similarity between AB and all non-corresponding BC subtracted as a baseline, within the time windows identified in the previous analysis (see **Figure 5B**). Bayesian linear regression was used to compare whether these neural similarities and dissimilarities show differences between trials with correct and incorrect indirect AC association or source memory. Both models controlled for repetition effects, and the source memory model also accounted for indirect AC association performance. To ensure balanced data, we excluded participants with near-ceiling indirect AC association or source memory performance (above 90%), and few incorrect responses. Thus, 30 participants were included in the model of AC association and 28 in the model of source memory.

We hypothesised that higher similarity between AB and corresponding BC movies, reflecting integration across related events, would be associated with higher AC retrieval. Conversely, greater dissimilarities, indicating efforts to separate related memory traces, would relate to better source memory. The observed results support our predictions. The similarity observed at 2.8 seconds was greater in trials with better AC memory performance (*β* = 0.044, BF_10_ = 14.668), suggesting that integrating events facilitates the formation of indirect associations. Additionally, the similarity observed at 3.7 seconds was negatively related to source memory performance (*β* = −0.036, BF_10_ = 5.345), indicating that integration may hinder the ability to remember specific event details. Finally, the dissimilarity observed at 7.4 seconds was associated with source memory performance (*β* = −0.041, BF_10_ = 8.681), showing that memory separation helps to preserve the source of different events (see **Figure 5C**). No other effects were significantly associated with AC or source memory performance (see **Supplementary Note 4** for more details).

### Electrophysiological Markers Associated with the Encoding of Related Events

A univariate analysis investigated the time-frequency patterns associated with encoding new events that share information with previously encoded experiences. A cluster-based permutation test^21^ was used to contrast BC encoding against XY encoding, within each BC movie segment and using the same frequency range (3-30 Hz) as in the previous representational similarity analysis (RSA). To be consistent with the previous RSA analysis, the initial encoding repetition of the BC and XY movies was excluded from this analysis.

Our results show increased alpha-beta power in the BC compared with the XY encoding in three different time windows: 1) between 0.0 to 1.5 seconds (cluster-*t* = −2.934e+4, *p* < .001), during the segment of *Sim C*; 2) between 2.8 to 4.5 seconds (cluster-*t* = −3.202e+4, *p* = .009), during the segment of *Sim C in Context*, and between 8.4 to 9.0 seconds (cluster-*t* = 2.505e+4, *p* = .014), during the segment of the *Sim B and C in Context*. Additionally, we observed a cluster indicating higher theta and alpha-beta power during the encoding of BC compared with XY between 6.4 to 8 seconds (cluster-*t* = −2.114e+4, *p* = .043), during the segment of *Sim B* (see **Figure 6**).

**Figure 6.**
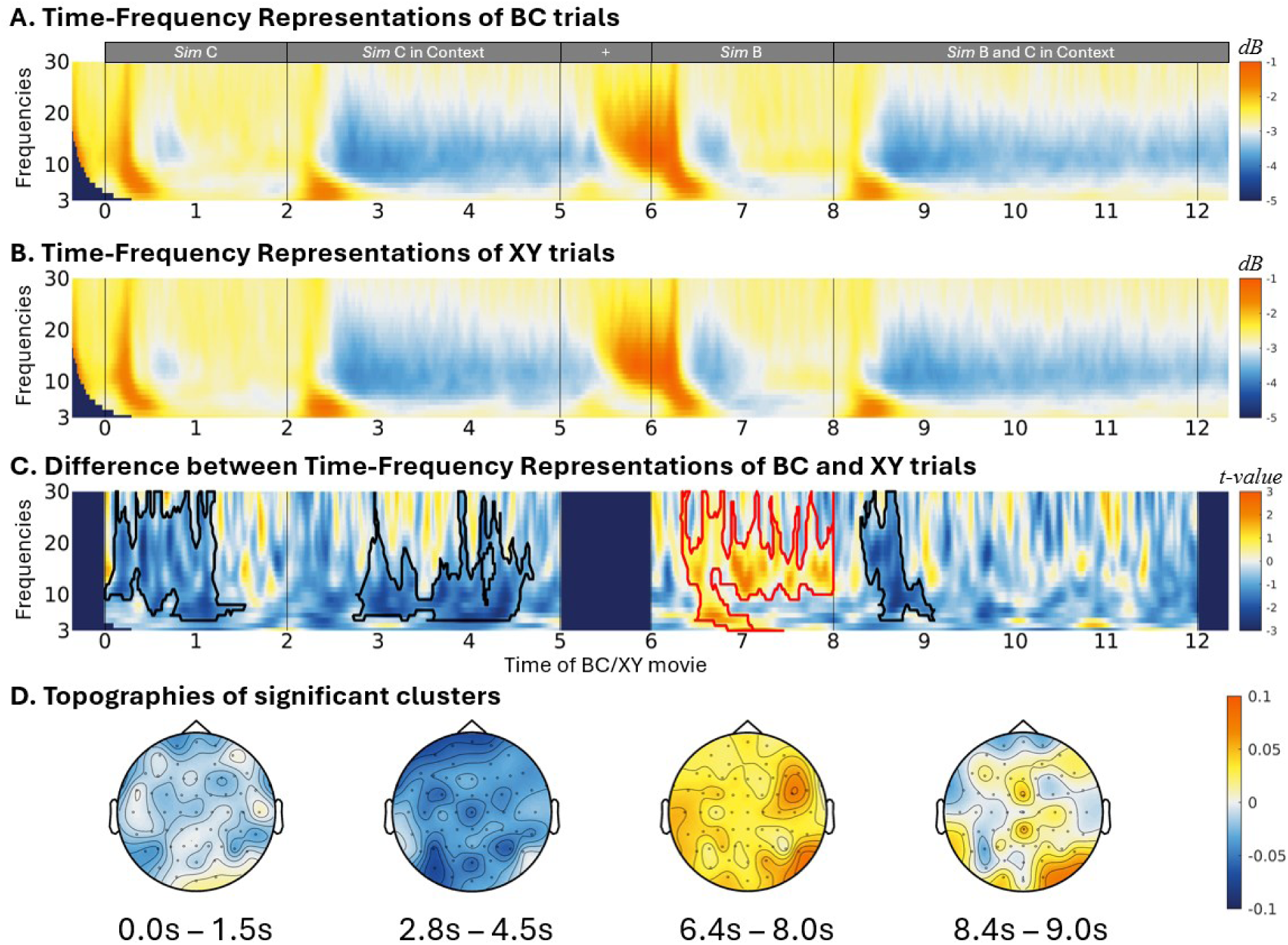
Time-frequency representations of BC (A) and XY (B) encoding, covering frequencies from 3 Hz to 30 Hz and their differences (C). The data were transformed into dB representations for visualization purposes. A cluster-based permutation analysis was used to estimate the power differences between BC and XY encoding. The significant clusters were marked in red (for BC > XY) and black (for BC < XY) edges. The topographies of the significant clusters of differences between BC and XY are plotted in (D).

Interestingly, the alpha-beta power decreases roughly correspond to the time windows where neural pattern similarities were found, while the increased theta and alpha/beta were found in the time windows associated with neural pattern dissimilarities. To examine if these time-frequency representations are linked to the formation of integrative and separated memory representation, we employed Bayesian linear regression analysis to investigate if these power changes predicted the neural (dis-)similarities observed in the corresponding time windows. The analysis showed that the alpha-beta power decrease observed between 0.0 to 1.5 seconds predicted the similarity centred at 1.0 second (*β* = −0.202, BF_10_ = 3.754e+29) during the segment of *Sim C*. Furthermore, the alpha-beta power decrease found between 2.8 seconds to 4.5 seconds predicted the similarity centred at 2.8 seconds (*β* = −0.045, BF_10_ = 29.831) and ∼3.7 seconds (*β* = −0.128, BF_10_ = 5.870e+11) during the segment of *Sim C in Context*. Additionally, the theta and alpha-beta power increase observed between 6.4 to 8 seconds predicted the dissimilarity centred at 7.4 seconds (*β* = −0.107, BF_10_ = 1.782e+8) during the segment of *Sim B*.

Previous studies have consistently reported that decreases in alpha-beta power are associated with successful memory encoding and retrieval^22–25^. Here, we provide evidence that alpha-beta power decreases are also related to the formation of integrated representations. Additionally, elevated alpha-beta power is typically linked to inhibitory control mechanisms^26^. Our findings suggest that increased alpha/beta power may be involved in establishing separated memory representations.

## DISCUSSION

The present study investigates how memories for related events are encoded into integrated and separated representations to support various mnemonic functions. Using an adapted version of the associative inference paradigm^5,16,17^ with naturalistic stimuli, Sim movies that simulate real-life interactions, we examined whether the brain flexibly encodes related events into integrated and separated memory representations. Participants first encoded a set of AB movies, followed by the encoding of a new set of BC movies, in which a familiar character interacted with a new one. A final memory test assessed participants’ ability to retrieve the indirect AC associations, while a source memory test examined if participants remembered that the two characters had not been seen together. By employing time-resolved representational similarity analysis (RSA) of EEG data, we tracked similarities and dissimilarities in the neural patterns between the encoding of the AB and BC movies. Our results show clear evidence that the human brain dynamically engages in memory integration and separation to support different aspects of memory, and provide novel insights into the temporal dynamics of these processes.

Increased representational similarity across related events has been consistently associated with forming integrated memory representations, which facilitates the linking of information from separate episodes into a coherent memory structure. This integrative process enables the extraction of commonalities across experiences, thereby supporting generalization and inference-making^6,7^. In contrast, representational dissimilarities have been implicated in maintaining distinctions between related events, thus reducing memory interference^1–3^. The present study expands these previous findings by demonstrating the simultaneous encoding of integrated and separated representations for related events (see **Figure 5**), each serving distinct memory demands. Specifically, we show that representational similarity enhances inferential memory performance by linking related events, whereas representational dissimilarity supports accurate source memory by preserving event-specific information. These findings refine our understanding of how the brain continuously modulates integration and separation in response to incoming information, offering insights into the neural mechanisms that balance flexibility and specificity in episodic memory formation.

The neural pattern similarities between AB and BC encoding were found related to successful AC retrieval (see **Figure 5**). These results are in line with the integrative encoding account^4,27^ and suggest that related events can be encoded into integrated representations that facilitate linked across event boundaries and support functions such as inferential reasoning and future decision making^6,7^. On the other hand, the neural pattern dissimilarities between AB and BC encoding was associated with better source memory performance (see **Figure 5**), indicating that memory separation efforts support the memory of specific event details^28,29^. Together, these highlight the idea that integration and separation mechanisms complement each other, allowing the brain to balance between forming generalized knowledge structures and maintaining distinct episodic memories. While integration aids inferential reasoning and decision by linking overlapping experiences, separation preserves the specificity of individual episodes and prevents interference.

Previous studies have reported that forming integrated memory representations impairs source memory and episodic details, implying that integrated and separated representations cannot coexist^10,11,18^. Here, we show that AC retrieval is associated with better source memory and found no evidence of a trade-off between memory integration and memory for specific events. Notably, however, the neural pattern similarities indicative of integration across AB and BC representations predicted reduced source memory accuracy (see **Figure 5**). This finding aligns with the idea that integration across events may blur episodic details, making it more difficult to recall the contents of a given event^10,11,18^. Our results suggest that the brain forms both integrated and separated representations in parallel. While the formation of integrated representation may lead to a loss of detail, this does not prevent the formation of a parallel representation for the individual details. The brain appears to simultaneously maintain integrated representations that facilitate indirect associations across related events, while retaining distinct representations for specific event details.

The use of EEG allowed us to capture the temporal dynamics of these memory processes. Interestingly, neural similarities between AB and BC encoding were predominantly observed in the time window corresponding to the presentation of the novel character C, suggesting that the presentation of novel information triggers its integration into an existing memory structure^6,7^. On the other hand, the neural dissimilarities between AB and BC encoding were observed when the previously seen character B was presented, suggesting that memory separation is triggered by the need of handling interference from overlapping information encountered across events^30,31^. Tracking these representations over repeated BC encoding trials revealed incremental shifts in similarity and dissimilarity patterns, indicating a gradual evolution of the memory representations (see **Supplementary Note 2**). This suggests that memory integration and separation are dynamic processes shaping the formation of memory representations across related experiences.

Using univariate analysis, we further identified the time-frequency representations (TFRs) associated with encoding new events that share information with past experiences. By contrasting the TFRs between BC and XY encoding, we found decreased alpha-beta power during the presentation of the novel C character and increased alpha-beta power during the presentation of the previously seen B character (see **Figure 6**). Interestingly, these alpha-beta power modulations roughly corresponded to the time-windows where the neural pattern similarities and dissimilarities between AB and BC encoding was found in the RSA. Specifically, alpha-beta power decreases were observed during the time-window with pattern similarities, while alpha-power increases occurred in the time-window with pattern dissimilarities. A Bayesian regression analysis confirmed that alpha-beta power decrease predicted neural pattern similarities, whereas the alpha-beta increase predicted neural pattern dissimilarities. Previous studies have consistently reported the involvement of alpha-beta power decreases in memory encoding and retrieval^22–25^, while alpha-beta power increases have been related with inhibitory control^26^. Our study suggests that alpha-beta power modulations are related to the formation of integrated and separated memory representations. The novel C character likely triggered the retrieval of the previously related AB event, leading to the integration of the C character into the existing memory representation. On the other hand, the increased alpha-beta power, observed when the shared character B was present, may have triggered inhibitory control, promoting the formation of separated memory representations.

In conclusion, by revealing the neural representational similarities and dissimilarities between related events, the present study demonstrates that integrated and separated representations can both be formed during learning and coexist to support different memory functions. Such learning across related events highlights the brain’s capacity for adaptive memory processing, which is crucial for making novel inferences while retaining the fidelity of individual episodes.

## METHOD

### Participants

We collected data from a total of 41 participants. Data from one participant was excluded for not having at least one incorrect AC association trial. Additionally, the data from another four participants were excluded due to poor EEG data quality and/or experimental programming errors. Hence, the final sample consisted of 36 participants (27 female, *M*_age_ ± *SD* = 24.3 ± 2.94).

Participants had normal or corrected-to-normal vision, were right-handed, fluent in English, and had no psychiatric or neurological disorders. Participants provided informed consent and were given a voucher that could be used in various shops for their time spent in the lab (on average 3 hours per participant including EEG preparations). The data collection was anonymous and did not involve any potentially identifying demographic information. The data collection was conducted in accordance with the Swedish Act concerning the Ethical Review of Research involving Humans (2003:460) and the Code of Ethics of the World Medical Association (Declaration of Helsinki). As established by Swedish authorities and specified in the Swedish Act concerning the Ethical Review of Research involving Humans (2003:460), the present study does not require specific ethical review by the Swedish Ethical Review Authority due to the following reasons: (1) it does not deal with sensitive personal data, (2) it does not use methods that involve a physical intervention, (3) it does not use methods that pose a risk of mental or physical harm, (4) it does not study biological material taken from a living or dead human that can be traced back to that person.

### Stimuli Material

AB and BC movies were generated using the life-simulation game *The Sims 4* by Electronic Arts (www.thesims4.com). In each movie, two Sims were presented. The movies were 14 seconds long and started with a fixation cross presented for 1 second, followed by the presentation of the Sim A/C for 2 seconds and the Sim A/C in a context, acting, for another 3 seconds. After another 1-second fixation cross, the Sim B was presented for 2 seconds followed by the Sim A/C interacting with Sim B in the same context for 5 seconds (see **Figure 2A**). Each movie has a unique context, so the AC association across AB and BC movies can only be made via the overlapping element, i.e., Sim B.

In total, 48 AB movies and 48 BC movies were created. Each AB movie has its own corresponding BC movie. During the experiment, only 24 AB movies were presented during the AB encoding, while all the BC movies were displayed during the BC encoding. As a result, participants could recognise the Sim B in 24 BC movies, while in the other 24 movies, the Sims were new to the participants. These all-new movies are the XY movies, which were used to control for memory processes involved in encoding direct associations without any previous overlapping experience. The selection and presentation of the AB movies were counterbalanced across participants.

In addition to the experimental movies, three videos of a man jogging at different places, also generated in *The Sims 4* environment, served as attention-check videos to keep participants concentrated.

To evaluate memory performance, 120 pictures of Sims faces (24 of each Sim A, B, C, X, and Y) were used as cues, targets, and distractors in the associative and source memory test. Additionally, to measure detail memory for context (see **Supplementary Note 1**), 96 pictures of contexts (48 for each AB and BC movie) were taken from the videos (without the Sim character present). One altered copy of each context was made by changing the colour and/or texture details of the original context, which resulted in 96 pictures of contexts with altered details. To test detail memory for clothing, 96 full-body pictures of Sims (48 of each Sim A and Sim C) were used. The distractor pictures for the clothing test were generated by changing the colour of the original clothing. For each Sim, three altered clothing pictures were generated, resulting in 288 pictures with altered details.

### Procedure

The stimuli were presented with PsychoPy^32^. The experiment consisted of an encoding and a test phase. In the encoding phase, participants first encoded 24 AB movies, each presented five times. When all 24 AB movies had been presented once, the next repetition began. After, participants encoded 24 BC movies intermingled with 24 XY movies, each presented also five times. The second repetition of these movies began when all of them had been presented one time. After the presentation of every 15 movies, the participants were encouraged to take a break. The attention-check movie was played three times during the encoding phase at random intervals. The participants were asked to press ENTER within three seconds each time they saw the attention-check movie. All included participants correctly responded to all attention checks within the given time.

The encoding phase ended with a short distraction task for 30 seconds, where participants were asked to consecutively subtract 7 from a random 3-digit number. Then, the test phase started. The test phase began with the direct and indirect memory tests, which were intermixed, with the only constraint that the AC indirect memory test was made before the corresponding AB and BC memory tests. Both the direct and the indirect associative memory test started with a cue, corresponding to the face of the Sim A or C, displayed on screen for 1.5 seconds, followed by the display of two faces presented below, the target Sim face C or A and the distractor Sim face drawn from the XY movies. Participants were asked to select which face was associated with the cue by pressing the left or the right key. Immediately after that, memory for source was tested. Participants were asked to indicate if the two Sims had been seen together in the same movie. For both association and source memory tests, participants were asked to rate their confidence on a three-point scale: 1 - guessing, 2 - maybe, 3 - sure. After the association and source memory tests, a surprise episodic detail memory test was also implemented to detect the memory integration-related episodic detail loss (see **Supplementary Note 1**).

Using Bayesian *t*-tests, we confirmed that indirect AC association performance was equivalent whether A or C was the cue (Accuracy: difference = 0.039, BF_01_ = 3.249; Response Time: difference = 0.027, BF_01_ = 4.099 and Confidence: difference = 0.002, BF_01_ = 4.113). The *BayesFactor* 0.9.12 package, based on R (4.1.2), was used to calculate the Bayesian Factors in favour of null hypothesis over the alternative hypothesis, BF_01_, with three as the significance criterion^33^. Prior gamma was set to 0.707 by default.

### Behavioural Data Analysis and Statistics

The behavioural data was analysed in R (4.1.2) using linear mixed models. The packages of *lme4* (1.1-34) and *emmeans* (1.8.7) were used to fit the models and perform statistical tests, and the *effectsize* (0.8.9) was used to estimate the effect sizes. The first level of each model was the trial level, which was clustered in the second level, i.e., the participant level. For all statistical analyses, the significance criterion of *p* < .05 was applied. Correct responses outside a 10 s time window or with a confidence rating of “guessing” were considered incorrect to prevent responses based on logical reasoning rather than memory to bias the results. Only correct responses were considered in the response times and confidence analysis.

First, behavioural performance for indirect and direct associative memory were contrasted in terms of accuracy, response times (RTs), and confidence ratings, with Association Type (AC vs AB vs BC vs XY) as the predictor for all three models. Next, source memory accuracy and confidence were also examined using Association Type (AB vs BC vs AC vs XY) and Association Accuracy (correct vs incorrect) of the preceding memory test as predictors. To interpret the interaction term observed in the source memory models, Monte Carlo sampling was implemented with in-house MATLAB scripts to estimate the effect size differences across all Association Types. For each Association Type, the effect sizes of the Association Accuracy over source memory accuracy and confidence were sampled for 10000 iterations, based on the model estimates obtained above. Then, these effect sizes were compared across Association Types by estimating the distribution of the difference between these samples. The percentage (%) of the sample difference that is larger/smaller than zero was considered an approximate of the one-tail statistical probability, which was then multiplied by two to indicate two-tail *p*-value, where the .05 criterion was applied to determine significance.

The normality of residuals was assessed by inspecting the Q-Q plots of the standardized residuals and Shapiro-Wilk’s tests. The homogeneity of variance of the model residuals was assessed for each model by visually inspecting a plot of the model residuals versus fitted values and using Levene’s test for unequal variance. Significant complex effects were followed up with post hoc tests with Tucky-correction. Effect sizes, partial eta squared (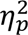) for F-tests and unstandardized difference (*D*) for t-tests, were also reported together with other statistics.

### EEG Data Collection and Preprocessing

EEG was recorded using a SynAmps RT Neuroscan 64-channel amplifier (sampling rate 1kHz, bandwidth DC-3500Hz, 24-bit resolution, left mastoid reference) with 62 electrodes attached to an elastic cap (active electrode EasyCap). The cap was placed according to the extended 10-20 system. Furthermore, an electrode was attached to the skin under the participants’ left eye to detect and later filter noise caused by eye blinks.

To preprocess the data, the EEG waveform was downsampled to 500Hz, and epochs of 14 seconds, ranging from −0.5 to 13.5 seconds with respect to the onset of each movie, were created. Preprocessing was implemented with FieldTrip^21^ accompanied by in-house scripts. The data were low-pass filtered at 200 Hz. A notch filter at 50, 100 and 150 Hz was applied to remove Alternate Current related signal. The data was transformed to linked-mastoid reference and demeaned. Vertical eye movements and blinks were estimated using the electrodes placed under the left eye and Fp1 in the cap, and horizontal eye movements were estimated using the Fp9 and Fp10 electrodes. The data were then manually checked to detect noisy channels and to remove trials dominated by artefacts other than eye blinks and horizontal eye movements. An independent component analysis was then applied to detect and remove components related to blinks and horizontal eye movements. Finally, after interpolating the removed channels, another visual check-up of the data took place for a final rejection of trials with residual artifacts. The cleaned data resulted, on average per participant, in 114 AB trials (ranging from 102 to 120), 114 BC trials (ranging from 100 to 119) and 114 XY trials (ranging from 107 to 119).

### Time-Frequency Decomposition of EEG data

The time-frequency decomposition was also implemented with in-house MATLAB scripts based on the Fieldtrip package^21^. The Morlet wavelets^34,35^ with 5 cycles were used to extract power spectra of frequencies of 3-30Hz (28 frequencies in total) for each event encoding trial (AB/BC/XY) with a temporal resolution of 0.02 second.

### Feature Selection for the Time-Resolved Representational Similarity Analysis

Before performing the representational similarity analysis, we aim to find the time-frequency features sensitive to AB movie content. This was done by estimating the wavelet coherence associated with encoding the AB movies. Wavelet coherence measures the correlation between the spectral components of two time-series^19^. By evaluating the coherence between all repetitions of the same AB movie and contrasting it against the coherence between different AB movies, we identified the time-frequency features sensitive to AB movie content. These features were later used to assess the representational similarities in the neural data between AB and BC/XY movies.

We calculated the pairwise wavelet coherence across all five repetitions of each AB movie for each participant, resulting in ten pairs per movie. As a control, we calculated the wavelet coherence between each AB movie and all other AB movies across all five repetitions, resulting in 230 pairs per movie. For each participant, the within-AB movie coherence was averaged across the 240 different pairs (24 different AB movies × 10 repetition pairs) and the between-AB movie coherence was averaged across 5470 pairs (24 different AB movies × 23 other AB movies × 10 repetition pairs).

A cluster-based permutation test was used to contrast the within-movie coherence (AB movie across repetitions) against the between-movie coherence (different AB movies) across participants. The test was conducted separately for four distinct time windows, corresponding to the different segments of the AB movie: 1) presentation of the Sim A, 2) presentation of Sim A in the context, 3) presentation of the Sim B, and 4) presentation of Sim A and B interacting in the context. For all comparisons, *t* values of significant clusters (two-tail *p* value ≤ .05), as well as their corresponding *p* values, are reported. Significant clusters indicated the time-frequency features sensitive to AB movie content, which were then used as templates to extract features (i.e., power at each channel-frequency-time) for the representational similarity analysis.

### Time-Resolved Representational Similarity Analysis and Statistics

The representational similarity between the neural patterns associated with AB encoding and the BC/XY encoding was estimated as a measure of the approximation and differentiation of mental representations^36^. As such, similarities in the neural representation are indicators of memory integration linking the events, while dissimilarities reveal memory separation between the events^9^.

The similarities between AB and BC/XY movies were estimated for each participant and movie. For the present analysis, the TFRs were normalised into decibel representation with respect to the average power of the whole epoch for each frequency. For each AB movie, a 3-dimensional representative time-frequency representation (channel-frequency-time) was obtained by averaging across the five repetitions. Representative features for each movie segment (*Sim A*, *Sim A in Context*, *Sim B*, and *Sim A and B in Context*) were extracted using the template obtained in the wavelet coherence analysis. Considering the time shifts across repetitions^37^, the medians along the time dimension of the extracted features were used as a stable estimation of movie AB. This procedure resulted in a 2-dimensional (channel-frequency) feature map for each AB movie segment, which was then reshaped into a feature vector by concatenating the values of each channel head to tail, representing the power distribution in the ‘channel-frequency’ space of a segment. In total, this feature extraction procedure resulted, for each participant, in 96 feature vectors (24 AB movies × 4 different AB movie segments) to be used in the similarity analysis contrasting the neural representations of AB and BC/XY movies.

The correlations between AB movies and BC/XY movies were performed separately for each of the five BC/XY repetitions, which enabled us to account for the progression of the similarities and dissimilarities along repetitions. The feature vectors corresponding to the AB movie contents were correlated with each time point of the BC/XY movie, resulting in a time-resolved representational similarity time series. For each BC/XY time point, the 2-dimensional matrix (channel-frequency) of power was reshaped in the same way as the extracted AB feature map, resulting in a vector representing the power distribution in the ‘channel-frequency’ space of each time point. This vector was correlated with the feature vector of each AB movie segment, resulting in 96 different correlation values (24 AB movies × 4 different segments), indicating the representational similarity between each segment of AB movie and each time point of each BC/XY movie. In the main paper we focus on the similarity towards the segment when the Sim A and B are interacting in a context, because this is the segment that comprises all the relevant information and encloses the AB movie event. The representational similarity analysis involving the other AB movie segments is presented in **Supplementary Note 3**.

To avoid the alpha error inflation induced by multiple comparison with conventional statistics, the significance of the similarity variation was determined by Bayesian statistics using an informative prior that accounts for false discoveries^38–40^, i.e., Normal-Inverse-Gamma distribution prior (alpha = 15, beta = 15, mu = 0, v = 30). The posterior was obtained by updating the prior with the data of all participants consecutively and tested against 0. A Bayesian Factor, in favour of alternative hypothesis over null hypothesis (BF_10_), larger than 3 is considered indicating significant difference^33^. Only the significant similarities and dissimilarities last over 0.1s were reported to avoid spurious results.

Note that when participants were first exposed to Sim C in the BC movies at repetition zero, we did not expect any memory reactivation of the AB movie. Thus, we excluded the repetition zero of the BC/XY trials from both the representational similarity analysis and the univariate analysis.

### Relationship between Behavioural Performance and Representational Similarities

To further reveal the behavioural consequences of representational similarities and dissimilarities between AB and BC, we investigated the relationship between the (dis-)similarities and behavioural performances in memory tests.

Bayesian linear regression with a Multivariate-Normal-Inverse-Gamma distribution prior (alpha = 15, beta = 15, mu = 0, lambda = 30 * *I, I* is unit matrix) and the data at trial level was used in the present analysis. The standardised regression coefficients were estimated to reduce the influences of individual differences. For each trial, the (dis-)similarities within each time window of significance were extracted and averaged as dependent variables. To estimate the (dis-)similarity difference between trials with correct and incorrect AC association, the accuracy of AC association was used as the independent variable, with the repetition order included as the controlled variable. Similarly, to estimate the (dis-)similarity difference between trials with correct and incorrect source memory for AC association, the accuracy of source memory was used as the independent variable, with the repetition order and the AC association accuracy included as the controlled variables. To ensure that the results were not biased by unbalanced data, the present analyses included only the participants with at least 10% incorrect trials for the independent variable (i.e., accuracy for AC association or source memory ≤ 0.9). Hence, the analysis for AC association included 30 participants, while the analysis for the source memory included 28 participants. A similar analysis for the clothing memory was also performed, see **Supplementary Note 4** for details.

### EEG Univariate Analysis and Statistics

A univariate analysis was used to investigate the neural correlates associated with the encoding of new events that share information with past events. To account for the individual difference in time frequency power during movie encoding, the difference between BC and XY was normalised by the power of BC for each participant, i.e., (BC-XY)/BC, and the statistical tests were performed by contrasting this index against zero. The comparison was performed separately for each segment of the movie (i.e., *Sim C* vs. *Sim Y*, *Sim C in Context* vs. *Sim Y in Context*, *Sim B* vs. *Sim X*, and *Sim C and B in Context* vs. *Sim Y and X in Context*) using a cluster-based permutation analysis implemented in fieldtrip^21^. For all comparisons, *t* values of significant clusters (two-tail *p* value ≤ .05), as well as their corresponding *p* values, are reported. Next, we evaluate if the potential differences between BC and XY encoding were predictive of the similarities and dissimilarities. This was performed using a Bayesian linear regression approach with a Multivariate-Normal-Inverse-Gamma distribution prior (alpha = 15, beta = 15, mu = 0, lambda = 30 * *I, I* is unit matrix) and the data at trial level. The power / similarity values were extracted from the trials according to the cluster / time-window of significance. Standardised regression coefficients were estimated to reduce influences of individual differences and the coefficient with BF_10_ > 3 was considered as significant effect^33^.

## Acknowledgements

This work was supported by the Swedish Research Council Grant VR 2019-02455. We thank Kira Friedrichs for the assistance in data collection and stimuli preparation and all the volunteers who participated in this study.

