## Supplementary Note for "Episodic events are flexibility encoded in both integrated and separated neural representations"

### Supplementary Note 1: Episodic Detail Memory Tests

At the end of the memory task, participants completed a surprise episodic detail memory test, assessing their memory for both context and the clothing of the characters A and C. For the context memory test, participants were first shown an image of the Sim for 1.5 s, followed by the presentation of four possible context pictures, arranged in a 2×2 matrix: 1) the original context of the Sim, 2) the context of Sim with altered details, 3) the context of the corresponding, indirectly associated Sim, and 4) the context of the corresponding, indirectly associated Sim with altered details. Participants were asked to select the context in which the Sim originally appeared (see **Figure S1A**). Following the context memory test, participants were asked to retrieve the colour and pattern of the Sim's clothing by choosing from four options (see **Figure S1B**). Participants rated their confidence on a three-point scale for all episodic detail memory tests: 1 – guessing, 2 – maybe, 3 – sure. There was no time limit for the responses, but responses made after 10s or associated with the confidence level 'guessing' were considered as incorrect response. For the current analyses, we included only trials where AB or BC associations were correctly retrieved, ensuring that surprise test results were based on accurate associative memory performance.

Memory for context was evaluated in two steps. First, we tested if participants incorrectly attributed the context of a Sim to its corresponding Sim due to memory integration. Previous studies suggest that, after forming an indirect AC association, participants may mistakenly recall the Sims' context as belonging to its corresponding Sim[1]. To this end, we ignored altered details and considered all trials correct for which participants selected the context of the current Sim (original or with altered details) and all trials incorrect for which the context of the corresponding Sim was selected (original or with altered details). A linear mixed-effects model tested if context memory varied as a function of AC association accuracy. The results showed no effect of AC accuracy, indicating no bias to select the context of the corresponding Sim when the AC association was correctly retrieved compared to when it was not ($F(1, 32) = 2.110$, $p = .156$, $\eta_p^2 = .06$).

Next, we tested whether participants retained the fine-grained contextual details following memory integration. We considered all trials correct for which participants selected the original, unaltered context and all trials incorrect for which participants selected the context with altered details. We separately analysed trials for which participants selected the contexts of the current Sim and trials for which participants selected the contexts of the corresponding Sim. Using linear mixed-effects models, we contrasted context memory as a function of AC accuracy (Correct vs. Incorrect). The results show that when participants selected the contexts of the current Sim, contextual detail memory performance was unaffected by AC accuracy ($F(1, 809) = 0.272$, $p = .602$, $\eta_p^2 = 3.36e\text{-}4$). The same was found for when participants selected the two contexts of the corresponding Sim ($F(1, 32) = 2.110$, $p = .156$, $\eta_p^2 = .06$). In summary, memory for contextual details was not affected by the AC memory performance (see **Figure S1C**).

For the clothing memory test, we examined whether its accuracy and confidence vary as a function of AC Accuracy. A marginal main effect of AC Accuracy was found for accuracy ($F(1, 1422) = 3.698$, $p = .055$, $\eta_p^2 = 2.59e\text{-}3$), but not for confidence ($F(1, 775) = 0.805$, $p = .370$, $\eta_p^2 = 1.04e\text{-}3$), suggesting that detail memory for clothing is likely worse after making correct AC association (.556) compared with an incorrect AC association (.612); see **Figure S1D**.

In sum, the results indicate that AC association performance did not systematically affect episodic memory details for context and clothing. Unlike source memory, which showed a clear link to AC retrieval, episodic detail memory appears to operate independently, suggesting that perceptual details may not be as strongly affected by the AC memory performance.

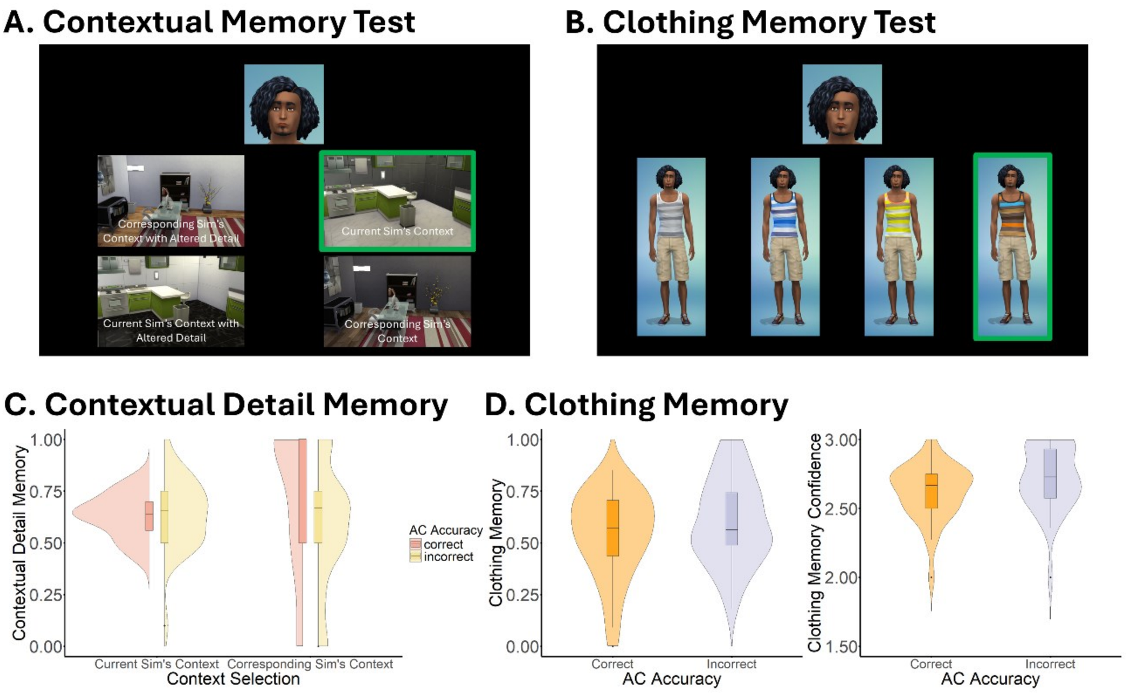


**Figure S1.** Surprise memory tests for episodic details and summary of the results. (A) Context memory was tested by asking participants to select the context associated with a given Sim from four different alternatives: 1) the current Sim's original context, 2) the current Sim's context with altered details, 3) the corresponding Sim's original context, and 4) the corresponding Sim's context with altered details. (B) Clothing memory was tested by asking participants to select the correct clothes of a given Sim. (C) Contextual detail memory accuracy: Proportion of trials where participants selected the original context over the altered version. Data is grouped by current Sim's context vs. corresponding Sim's context and further split by AC association accuracy. (D) Clothing memory accuracy and confidence as a function of indirect AC association accuracy.

### Supplementary Note 2: Representational Similarity Across the Five Repetitions of Each Movie

Here, we show how the neural similarities and dissimilarity evolve across the five encoding repetitions of the BC movie (**Figure S2**). For each participant, movie, and repetition, the neural (dis-)similarity values were extracted for the time windows with significant results in the main analysis (see **Figure 4**).

First, we tested if these neural (dis-)similarities were already present in the initial encoding of the BC movie by contrasting the patterns of the initial repetition against zero. This hypothesis was not supported by the data ($BF_{10}$s < 0.998), suggesting that no significant similarities or dissimilarities were detected during the initial repetition. This is not surprising because, during the first repetition, participants would not yet know the correspondence between AB and BC movies until Sim B was presented.

Next, we tested whether a linear trend emerged for neural (dis-)similarities across the five repetitions. A Bayesian linear regression was applied to predict the neural (dis-)similarities throughout the five rounds of encoding. The results showed that the similarities observed at 1.0 second followed such a trend during the first three encoding rounds ($\beta = 0.138$, $BF_{10} = 3.789$). The linear progressive pattern was also observed for the similarity observed at 3.7 seconds ($\beta = 0.125$, $BF_{10} = 5.291$) across the five encoding rounds. However, no significant linear trend was found for the similarity observed 2.8 seconds ($BF_{10}$s < 1.693). The dissimilarity observed at 7.4 seconds ($\beta = -0.157$, $BF_{10} = 8.809$) showed a linear trend for the first four rounds.

These findings suggest that neural similarities and dissimilarities gradually shift over the five encoding repetitions. The observed linear trends in similarity and dissimilarity likely indicate an iterative adaptation of the integrated and separated representations over time, as the brain progressively refines the memory encoding process with each repetition.

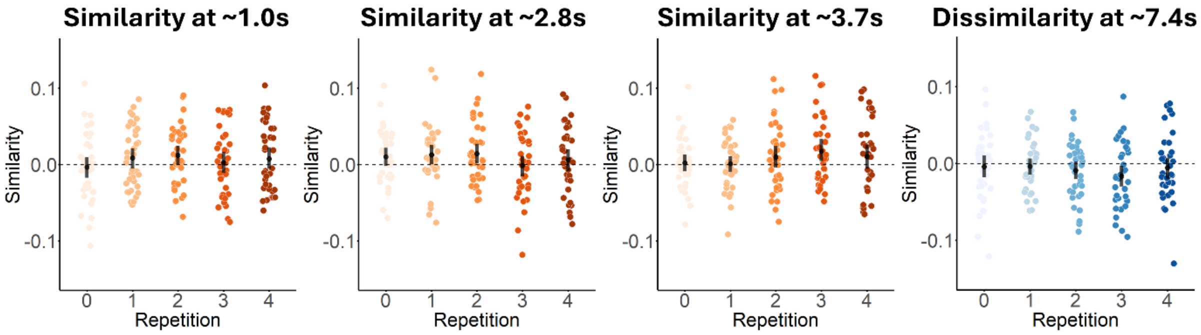


**Figure S2** Progressions of similarities and dissimilarity across the five encoding repetitions of the BC movie. The 95% CI with bootstrapping is plotted for reference. Each dot represents the data of a participant. Similarities are indicated in orange and the dissimilarity in blue, the depths of which refer to different repetitions.

**Supplementary Note 3: Representational Similarity Analysis between Other AB Segments and Corresponding BC Movies**

Beyond assessing the representational similarity between the segment corresponding to when Sim A and B interacted in a Context and the corresponding BC movie, we also estimated the similarity between other segments of AB movie and their corresponding BC movie. This analysis aimed to provide a more comprehensive view of how different phases of AB encoding relate to subsequent BC representations. The temporal dynamics of these similarities are plotted in **Figure S3**, highlighting fluctuations in the neural representational across different time windows. The specific time windows of significant similarity, along with their statistics, are summarised in **Table S3**.

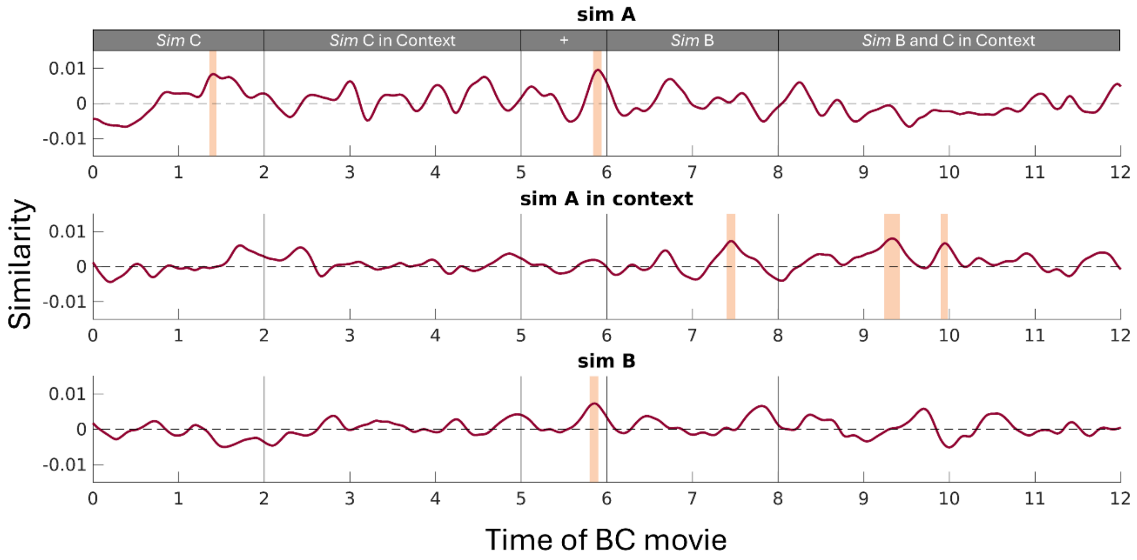


**Figure S3.** Neural representational similarities and dissimilarities between other AB movie segments and BC movie encoding. The times corresponding to significant differences are marked in highlighted and the statistics are summarized in **Table S3**.

**Table S3.** Summary of the significant time windows and statistics between other segments of AB movie and BC movie encoding.

| | Time | Average Similarity | $BF_{10}$s > |
|---|---|---|---|
| *Sim A* | | | |
| | 1.4s | 8.4e-3 | 3.063 |
| | 5.9s | 9.3e-3 | 3.160 |
| *Sim A in Context* | | | |
| | 7.5s | 7.1e-3 | 3.099 |
| | 9.3s | 7.5e-3 | 3.410 |
| | 10.0s | 6.6e-3 | 3.003 |
| *Sim B* | | | |
| | 5.9s | 7.2e-3 | 3.408 |

**Supplementary Note 4: Relationship between Neural Pattern Similarities / Dissimilarities and Behavioural Memory Tests**

Here we report the overall relationships between all the systematic (dis-)similarities and the behavioural performance of the memory tests, including AC inference, source memory, and memory for clothing. Memory for clothing was also included in the present analysis because it likely covaries with AC memory performance (see **Supplementary Note 1**). Specifically, we estimated how the similarity values relate to later clothing memory, with the repetition order and AC inference accuracy included as controlled variables. The results are summarised in **Table S4**.

In addition to what is reported in the main text, we observed that higher neural pattern similarity between Sim A observed at 1.4 seconds during BC movie encoding was associated with later AC inference performance. This aligns with our main finding, showing that the neural pattern similarities reflect the formation of an integrated representation involving the elements encountered across different events, which can later be used to respond to the AC memory test. Additionally, the similarities between Sim A in Context and BC movie, observed at 9.3 seconds and 10.0 seconds, were negatively related to memory for clothing, indicating that the formation of an integrated representation may lead to the loss of episodic details associated with the individual events.

**Table S4.** Bayesian Linear Regression-based analysis investigating the relationship between the representational (dis-)similarities and the behavioural memory tests. Significant effects are marked in bold.

| Segment | Time | AC inference accuracy | | Source memory accuracy | | Clothing memory accuracy | |
|---|---|---|---|---|---|---|---|
| | | *β* | $BF_{10}$ | *β* | $BF_{10}$ | *β* | $BF_{10}$ |
| Similarity to *Sim A* | | | | | | | |
| | 1.4s | **0.049** | **26.268** | -0.014 | 1.283 | -0.031 | 2.755 |
| | 5.9s | 0.012 | 1.214 | 0.028 | 2.811 | 0.005 | 1.013 |
| Similarity to *Sim A in Context* | | | | | | | |
| | 7.5s | -0.027 | 2.779 | -0.003 | 1.004 | 0.013 | 1.168 |
| | 9.3s | -0.004 | 1.011 | -0.011 | 1.154 | **-0.033** | **3.021** |
| | 10.0s | -0.006 | 1.046 | 0.001 | 0.992 | **-0.044** | **7.756** |
| Similarity to *Sim B* | | | | | | | |
| | 5.9s | -0.002 | 0.996 | 0.013 | 1.238 | 0.005 | 1.016 |
| Similarity to *Sim A and B in Context* | | | | | | | |
| | 1.0s | 0.022 | 1.922 | 0.000 | 0.992 | -0.013 | 1.169 |
| | 2.8s | **0.044** | **14.668** | 0.022 | 1.820 | 0.025 | 1.931 |
| | 3.7s | 0.000 | 0.992 | **-0.036** | **5.345** | -0.004 | 1.009 |
| | 7.4s | 0.012 | 1.222 | **-0.041** | **8.681** | 0.001 | 0.992 |
| *N* of participant | | | 30 | | 28 | | 36 |
